# Central deficiency of norepinephrine synthesis and norepinephrinergic neurotransmission contributes to seizure-induced respiratory arrest

**DOI:** 10.1101/865691

**Authors:** Yue Shen, HaiXiang Ma, Han Lu, HaiTing Zhao, Jianliang Sun, Yuan Cheng, Yi Shen, Yu Dong Zhou, HongHai Zhang

## Abstract

**Objective:** Sudden unexpected death in epilepsy (SUDEP) is the leading cause of mortality in patients in patients with intractable epilepsy. However, the pathogenesis of SUDEP seems to be poorly understood. Our previous findings showed that the incidence of seizure-induced respiratory arrest (S-IRA) was markedly reduced by atomoxetine in a murine SUDEP model. Because the central NE α-1 receptor (NEα-1R) plays a vital role in regulating respiratory function, we hypothesized that the suppression of S-IRA by atomoxetine was mediated by NE/NEα-1R interactions that can be reversed by NEα-1R antagonism.

**Methods:** We examined whether atomoxetine-mediated suppression of S-IRA evoked by either acoustic stimulation or pentylenetetrazole (PTZ) in DBA/1 mice can be reversed by intraperitoneal (IP) and intracerebroventricular (ICV) administration of prazosin, a selective antagonist of NEα-1R. The content and activity of tyrosine hydroxylase (TH), a rate-limiting enzyme for NE synthesis, in the lower brainstem was measured by ELISA. Electroencephalograms (EEG) were obtained by using the PTZ-evoked SUDEP model.

**Results:** Atomoxetine-mediated suppression of S-IRA evoked by either acoustic stimulation or PTZ was significantly reversed by low doses of IP and ICV prazosin. Neither repetitive acoustic stimulation nor S-IRA reduced TH levels in lower brainstem. However, the enzyme activity of TH levels in lower brainstem was significantly increased by mechanical ventilation with DBA/1 mice ,which makes dead DBA/1 mice suffered from S-IRA and SUDEP recover. EEG data showed that although the protective effect of atomoxetine was reversed by prazosin, neither drug affected EEG activity.

**Significance:** These data suggest that deficient synthesis of NE and norepinephrinergic neurotransmission contributes to S-IRA and that the NEα-1R is a potential therapeutic target for the prevention of SUDEP.

## 1. INTRODUCTION

Sudden unexpected death in epilepsy (SUDEP) had been recognized as the most cause for death of epilepsy patients, which accounts for up to 17% of deaths in epilepsy patients, in the case of potential life lost from SUDEP, ranking second only to the stroke (Bateman,et al., 2010; Tomson, et al., 2008; Lhatoo, et al., 2015). Although the pathogenesis of SUDEP is unclear, multiple studies indicate that most cases are caused by cardiopulmonary dysfunction (Devinsky,et al., 2016). Our previous findings demonstrated that seizure-induced respiratory arrest (S-IRA) is the most important cause of SUDEP evoked by either acoustic stimulation or pentylenetetrazole (PTZ) in DBA/1 murine models, and was significantly reduced by atomoxetine, a selective norepinephrine (NE) reuptake inhibitor that is commonly used for treating attention-deficit hyperactivity disorder (ADHD) (Zhang,et al., 2016; Gordon and George, 2016; Zhang,et al., 2017; Zhang,et al., 2017; Zhang, et al.,2018). However, the mechanisms underlying atomoxetine suppressing S-IRA are unknown (Devinsky,et al., 2016). In addition, our previous studies that used multiple epilepsy models showed that central excitatory synapses promote the pathogenesis of epilepsy (Zhou,et al., 2009; Shen, et al.,2018). Because the NE α-1 receptor (NEα-1R) plays a vital role in the central regulation of respiration (Doi A,et al., 2010; Zanella, et al.,2014; Schnell et al.,2018; Zanella, et al.,2016), we hypothesized that it may be a key therapeutic target to prevent SUDEP.

To study the roles of NE and NEα-1R in the pathogenesis of S-IRA in DBA/1 mice from behavioral and electrophysiologic perspectives, we first attempted to enhance NE bioavailability in the central synaptic spaces by using atomoxetine. We then sought to determine whether the atomoxetine-mediated suppression of S-IRA can be reversed by the peripheral and central administration of prazosin, a selective NEα-1R antagonist. To exclude the possibility that neurologic effects of prazosin could be mediated by altered cardiovascular function, we measured changes of mean systolic blood pressure (SBP), diastolic blood pressure (DBP), mean arterial pressure (MAP), and heart rate (HR) in DBA/1 mice in different dosage groups. Furthermore, we studied the effect of prazosin on atomoxetine-mediated suppression of S-IRA by evaluating seizure scores and electroencephalographic (EEG) recordings.

In our study, atomoxetine-mediated suppression of S-IRA following acoustic stimulation and PTZ injection in DBA/1 mice was significantly reversed by intraperitoneal (IP) and intracerebroventricular (ICV) injections of prazosin in two models, without affecting seizure behavior, cardiovascular function, or EEG activity. In addition, our results provide the first evidence that the occurrence of S-IRA were attributed to the decreased activity of TH in lower brainstem, which led to the lower synthesis of NE in brainstem. Thus, our results suggest that enhancing central the synthesis of NE and norepinephrinergic neurotransmission, and targeting the central NEα-1R may represent rational therapeutic strategies to prevent SUDEP.

## 2. MATERIALS AND METHODS

### 2.1 Animals

All experimental procedures were compliant with the National Institutes of Health Guidelines for the Care and Use of Laboratory Animals and approved by the Animal Advisory Committee at Zhejiang University. DBA/1 mice were housed and bred in the Animal Center of Zhejiang University School of Medicine and provided with rodent food and water ad libitum. In the acoustic stimulation murine model, DBA/1 mice were “primed” starting from postnatal day 26-28 to establish consistent susceptibility to audiogenic seizures and S-IRA. Atomoxetine (dissolved in saline and given IP) and prazosin were administered 120 mins and 30 mins prior to acoustic stimulation, respectively. In the another model of PTZ-evoked seizure model, PTZ was administered to non-primed DBA/1 mice of approximately 8 weeks of age. The two models were established just as described previously (Zhang, et al., 2016; Zhang,et al., 2017; Zhao,et al., 2017; Zhang, et al.,2018).

### 2.2 Seizure induction and resuscitation

S-IRA was evoked by acoustic stimulation and intraperitoneal (IP) administration of PTZ, as previously described (Zhang,et al., 2016; Zhang, et al., 2017; Zhao, et al., 2017; Zhang, et al., 2018). For the acoustic stimulation model, each mouse was placed in a cylindrical plexiglass chamber in a sound-isolated room, and audiogenic seizures (AGSZs) were evoked by an electric bell (96 dB SPL, Zhejiang People’s Electronics, China). Acoustic stimulation was given for a maximum duration of 60 s or until the onset of tonic seizures and S-IRA in most mice in each group. Mice with S-IRA were resuscitated within 5 s post the final respiratory gasp using a rodent respirator (YuYan Instruments, ShangHai, China). S-IRA was evoked in all non-primed DBA/1 mice by the IP administration of a single dose of PTZ (Cat # P6500; Sigma-Aldrich, St. Louis, MO) at a dose of 75 mg/kg. Mice with S-IRA were resuscitated by using a small animal respirator (YuYan Company, ShangHai, China).

### 2.3 Effect of IP prazosin on atomoxetine-mediated suppression of S-IRA

A vehicle control group and treatment groups using different doses of prazosin (Cat # P7791; Sigma-Aldrich) were evaluated using the acoustic stimulation model using atomoxetine (Ca # Y0001586; Sigma-Aldrich) pre-treatment. Susceptibility to S-IRA in primed DBA/1 mice was confirmed 24 h before atomoxetine or vehicle and prazosin administration. Atomoxetine (15 mg/kg) and prazosin (0.001-0.1mg/kg) were given by IP injection 120 min and 30 min prior to acoustic stimulation, respectively. In the vehicle control group, saline and 25% dimethyl sulfoxide (DMSO) were given by IP injection 120 min and 30 min prior to acoustic stimulation respectively. The incidence of S-IRA latency to AGSZs, duration of wild running, clonic seizures, tonic-clonic seizures, and seizure scores were videotaped for offline analysis (n=6-9/group) (Zhang, et al., 2016; Zhang, et al., 2017; Zhao, et al., 2017; Zhang, et al., 2018).

### 2.4 Effect of IP prazosin on S-IRA

A vehicle control group and treatment groups given different doses of prazosin (Cat # P7791; Sigma-Aldrich) were evaluated using included in the acoustic stimulation model. Susceptibility to S-IRA was confirmed 24 h prior to the administration of vehicle and prazosin. In the vehicle control group, saline and 25% DMSO were given by IP injection 120 min 30 min prior to acoustic stimulation, respectively. In the treatment groups, saline and prazosin (0.005-2mg/kg, dissolved in 25% DMSO) were administered by IP injection 120 min and 30 min prior to acoustic stimulation, respectively. Incidence of S-IRA, latency to AGSZs, duration of wild running plus clonic seizures (W+C), tonic-clonic seizures and seizure scores were videotaped for offline analysis (Donato, et al., 2007; Soper, et al., 2016). (n = 6-9/group)

### 2.5 Effect of IP prazosin on blood pressures and heart rate

Non-invasive measurements of heart rate and blood pressures at 37°C were performed in conscious DBA/1 mice treated with either atomoxetine or vehicle using a tail-cuff system (BP-98A, Blood Pressure Analysis System, Softron, Japan). In the control group, atomoxetine (15 mg/kg) and 25% DMSO were given by IP injection 120 min and 30 min prior to acoustic stimulation, respectively. The experimental group was given atomoxetine (15 mg/kg) and prazosin (2mg/kg dissolved in 25% DMSO) by IP injection 120 min and 30 min prior to acoustic stimulation, respectively. Measurements of SBP, DBP, MAP and HR were taken during 60 s intervals at 120 min and 25 min before and 5 min after acoustic stimulation.

### 2.6 Effect of ICV prazosin on S-IRA

An ICV guide cannula (O.D.I.41×I.D.O.0.25mm/M3.5, 62004, RWD Life Science Inc.,China) was implanted into the right lateral ventricle (AP – 0.45 mm; ML – 1.0 mm; V– 2.50 mm) to enable microinjections as previously described (Zhao, et al., 2017). The experimental groupings were: 1) Saline (IP) was administered 120 min prior to PTZ (75mg/kg, IP) and 25% DMSO at (2 μl, at a rate of 0.5 μl/min ICV) 15 min prior to PTZ injection as controls. 2) Atomoxetine (15mg/kg, IP) was administered 120 min prior to PTZ (75mg/kg, IP) and 25% DMSO (2 μl, at a rate of 0.5 μl/min ICV) 15 min prior to PTZ injection. 3) Atomoxetine (15mg/kg, IP) was administered 120 min prior to PTZ (75mg/kg, IP), and prazosin (4.764 nmol and 9.528 nmol, dissolved in 2 μl 25% DMSO, at a rate of 0.5 μl/min ICV) 15 min prior to PTZ injection (n=6-9/group).

### 2.7 Effect of ICV prazosin on EEG

After the implantation of ICV guide cannulas described above, DBA/1 mice were implanted with a headstage for EEG recordings. The headstage consisted of a 6-pin connector soldered to 2 EEG screws. The EEG electrode was inserted into the surface of cerebral cortex fixed by screws inserted into the cranium with a layer of dental cement mixed with cyanoacrylate glue. EEG recordings were performed one week after surgery. The experimental groups (n=6-9/group) were: 1) Saline (IP) and 25% DMSO (2 μl, at a rate of 0.5 μl/min ICV) at 120 min and 15 min prior to PTZ (75mg/kg, IP), respectively, as controls. 2) Atomoxetine (15mg/kg, IP) and 25% DMSO (2 μl, at a rate of 0.5 μl/min ICV) 120 min and 15 min prior to PTZ (75mg/kg, IP), respectively. 3) Atomoxetine (15mg/kg, IP) and prazosin (4.764 and 9.528 nmol dissolved in 2 μl 25% DMSO administered a rate of 0.5 μl/min ICV) 120 min and 15 min prior to PTZ (75mg/kg, IP), respectively. EEG recording began immediately to after the completion of ICV injections. ICV placement was confirmed by histology with DAPI staining as previously described (Zhao, et al., 2017).

### 2.8 Measurement the contnet and activity of tyrosine hydroxylase (TH) in brianstem and prosencephalon by ELISA

DBA/1 mice from 3 groups that included a naive group without acoustic stimulation, an S-IRA evoked by acoustic stimulation group, and an S-IRA evoked by acoustic stimulation with resuscitation by small animal respirator group were decapitated under anesthesia using an overdose of chloral hydrate (n = 6/group). As we described previously (Zhang, et al., 2016), the brainstems between bregma −3 mm and bregma −9 mm and the prosencephalon were collected after removing excess blood in cold PBS and then immediately frozen in liquid nitrogen and transferred to a −80°C refrigerator. Tissue specimens were weighed and then homogenized in ice-cold PBS. Homogenates were centrifuged for 5 minutes at 5000g to obtain supernatant. Both content and enzyme activity of TH was evaluated by ELISA, respectively. All ELISA procedures were performed following the manufacturer’s instructions (Mouse TH, ELISA Kit, YanSheng Biological Technology Co., Ltd, ShangHai, CS1028; CS03271, China).

## 3.0 STATISTICAL ANALYSIS

All data are presented as the mean ± SEM. Statistical analyses were performed using Prism 8.0 software (GraphPad Software, La Jolla, CA, USA) .The incidence of S-IRA in different groups was compared using Wilcoxon Signed Rank test. Seizure scores and the latency to AGSZs, the duration of wild running, clonic seizures, tonic-clonic seizures, content and activity of TH, average EEG spectra were evaluated using the one-way ANOVA tests. Statistical significance was inferred if p < 0.05.

## 4. RESULTS

### 4.1 Atomoxetine-mediated suppression of S-IRA evoked by acoustic stimulation was reversed by prazosin

Compared to vehicle controls, the incidence of S-IRA evoked by acoustic stimulation was significantly reduced by atomoxetine at an IP dose of 15 mg/kg (p<0.01). Atomoxetine-mediated suppression of S-IRA by was significantly reversed by IP prazosin (p<0.01). There were no significant intergroup differences in latencies to AGSZs, durations of wild running, clonic seizures, tonic-clonic seizures and seizure scores (p > 0.05) (Fig. 1).

**Figure 1.**
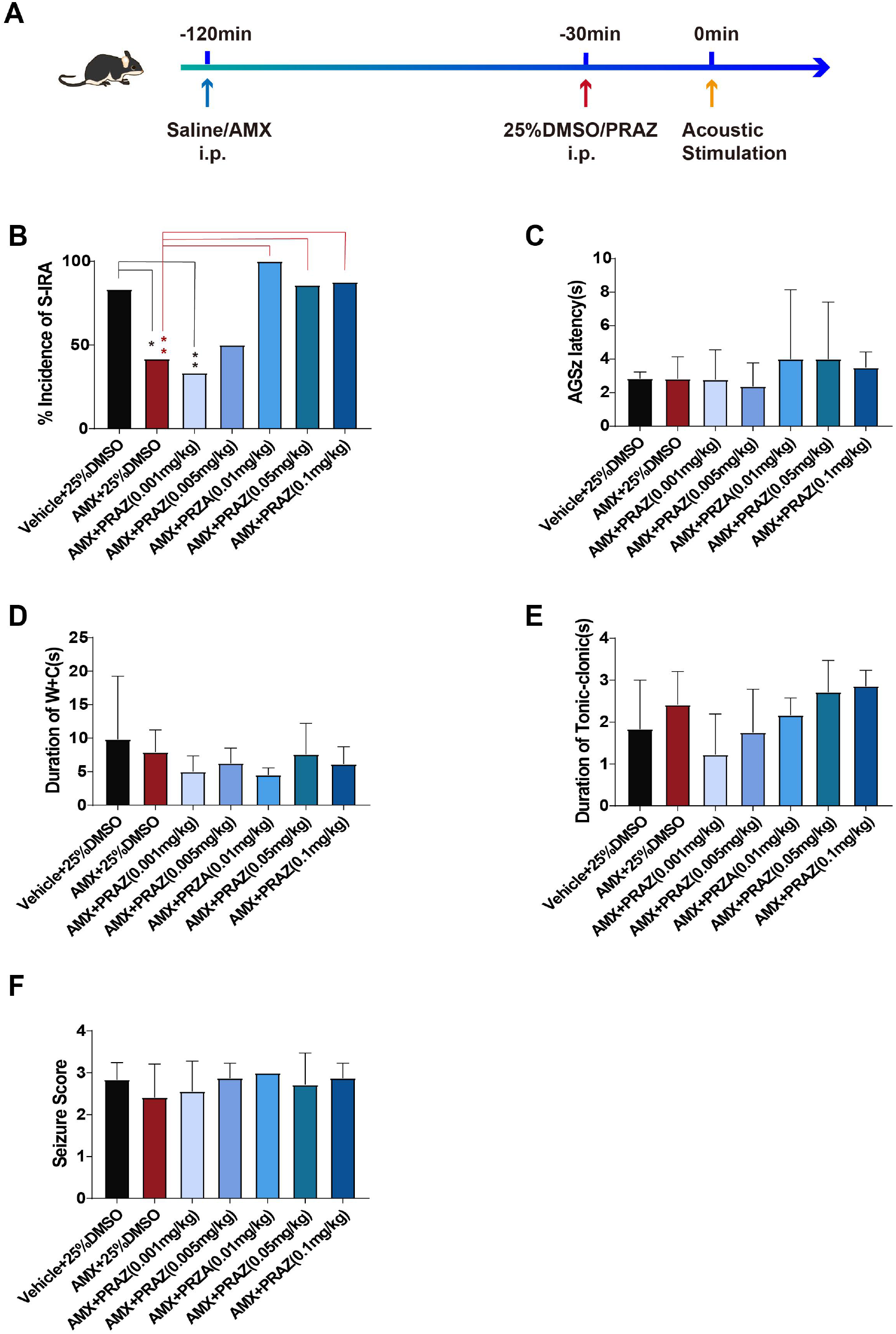
Atomoxetine-mediated reduction of S-IRA evoked by acoustic stimulation was significantly reversed by prazosin. B. Compared to the control group (n=6), S-IRA evoked by acoustic stimulation was markedly lower in groups treated with i.p. atomoxetine (AMX) (n=12) or AMX and prazosin (PRAZ) at 0.01-0.05mg/kg (n=9,8) in primed DBA/1 mice (** p < 0.01). However, the protective effect of AMX in primed DBA/1 mice was significantly reversed by prazosin doses of 0.01-0.1mg/kg (n=9,8,6,7,8) (** p < 0.01). C-F. There were no intergroup differences in latencies to AGSZs, durations of wild running plus clonic seizures (W+C), tonic-clonic seizures, and seizure scores (p > 0.05). AMX was administered 120 min and PRAZ was given 30 min prior to induction of AGSZs. S-IRA, seizure-induced respiratory arrest; AGSZs, audiogenic seizures; i.p., intraperitoneal injection; DMSO, dimethyl sulfoxide.

### 4.2 The decreased activity of TH resulted in S-IRA was significantly reversed by mechanical ventilation in lower brainstem

We examined the content and activity of TH in the brainstem of DBA/1 mice from 3 groups that included a control group without acoustic stimulation, and two S-IRA groups following acoustic stimulation with and without resuscitation, respectively. Compared to the relative optical density of DBA/1 mice without acoustic stimulation, to which the other two groups were normalized, TH content was not significantly reduced in the brainstem of primed DBA/1 mice (p > 0.05, n=6). TH levels were not significantly different between the S-IRA with and without resuscitation groups (p > 0.05, n=6). However, compared with treatment of brainstem, there was a significant reduction of TH content in prosencephalon in the control and S-IRA (rescue) groups. (Fig. 2). Compared with control treatment of brainstem, TH activity was significantly decreased in prosencephalon in control, S-IRA and S-IRA (rescue) groups.(p < 0.01, n=6). Compared with the TH activity in brainstem in control group, there was a significant reduction in brainstem in S-IRA (rescue) group. (p < 0.001, n=6). Compared with the TH activity in prosencephalon in control group, there was a significant reduction in prosencephalon in S-IRA (rescue) group. (p < 0.001, n=6). (Fig. 2).

**Figure 2.**
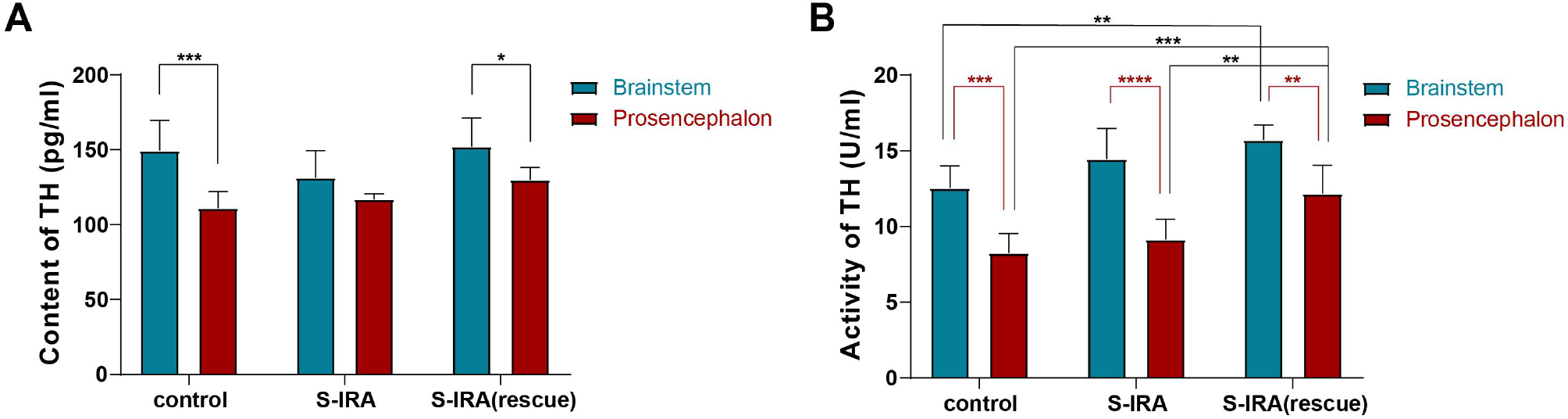
Continuous *acoustic stimulation did not change the content of TH but change its activity in brainstem*. Compared to the control group without acoustic stimulation, there were no significant changes in TH levels in lower brainstem in the S-IRA without resuscitation and S-IRA with resuscitation groups (n=6) (p > 0.05). There were no significant differences of TH levels in lower brainstem between S-IRA without resuscitation and S-IRA with resuscitation groups (n=6) (p > 0.05). However, the content of TH in lower brainstem was significantly than in prosencephalon in control and S-IRA (rescue) groups. Compared with the TH activity in brainstem in control group, there was a significant reduction in brainstem in S-IRA (rescue) group. (p < 0.001, n=6). Compared with the TH activity in prosencephalon in control group, there was a significant reduction in prosencephalon in S-IRA (rescue) group. (p < 0.001, n=6).

### 4.3 Effects of prazosin on blood pressure and heart rate

Given that prazosin can lower peripheral blood pressures, affect heart rate, and reduce cerebral perfusion pressure to cause S-IRA in our model, we measured changes of SBP, DBP, MAP and HR 5 min after acoustic stimulation. There were no significant differences of SBP, DBP, MAP and HR between the vehicle control group in which IP atomoxetine (15 mg/kg) and 25% DMSO were given 120 min and 30 min prior to acoustic stimulation, respectively; and the experimental group (n = 6) in which IP atomoxetine (15 mg/kg) and prazosin were given 120 min and 30 min prior to acoustic stimulation, respectively (Fig. 3).

**Figure 3.**
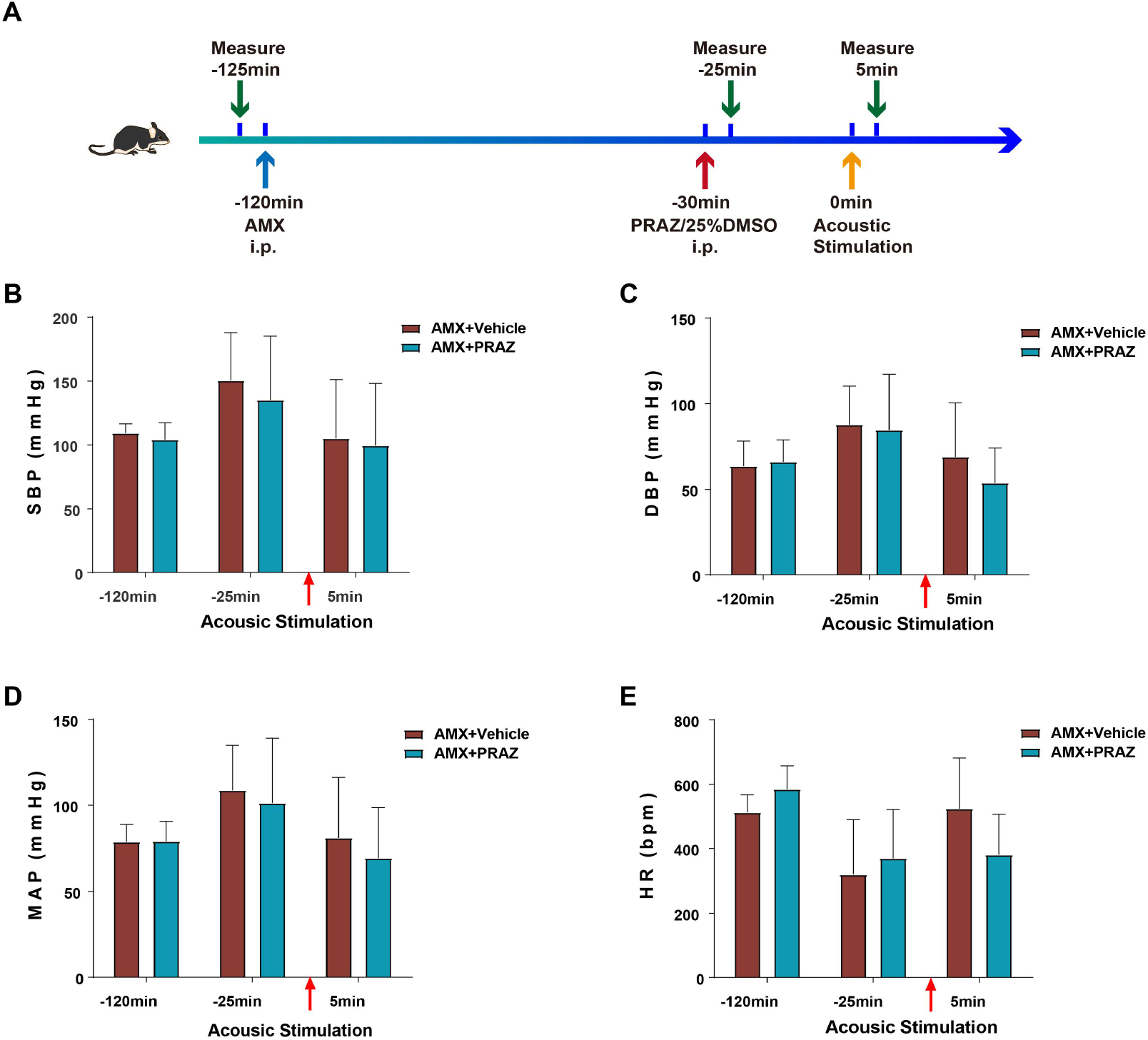
Effects of prazosin on the blood pressure and heart rate. B-E. Compared to the control group (n=6) in which atomoxetine (AMX) was given i.p. 120 min and 25% DMSO given i.p. 30 min before and 5 min after acoustic stimulation, there were no significant differences in SBP, DBP, MAP and HR in the group in which AMX was given 120 min prior to acoustic stimulation, and prazosin (PRAZ i.p) (n=6) 30 min before and 5 min after acoustic stimulation (p >0.05). SBP, systolic blood pressure; DBP, diastolic blood pressure; MAP, mean arterial pressure; HR, heart rate; i.p., intraperitoneal.

### 4.4 Effects of ICV prazosin on atomoxetine-mediated suppression of S-IRA evoked by PTZ

Because of possible differential effects of prazosin on atomoxetine-mediated suppression of S-IRA evoked by different epileptogenic stimuli, we chose an established SUDEP model using PTZ to induce S-IRA. Compared to the vehicle control group, PTZ-induced S-IRA was markedly suppressed by atomoxetine (p< 0.01). Compared to the control group (IP atomoxetine, 15mg/kg, ICV DMSO, n=6), low-dose prazosin (atomoxetine, 15mg/kg, i.p, prazosin, 4.764 nmol,icv) did not reverse the atomoxetine-mediated suppression of S-IRA (p>0.05). However, atomoxetine-mediated suppression of S-IRA by was significantly reversed by a higher prazosin dose (9.528 nmol, icv) (p < 0.01) (Fig. 4).

**Figure 4.**
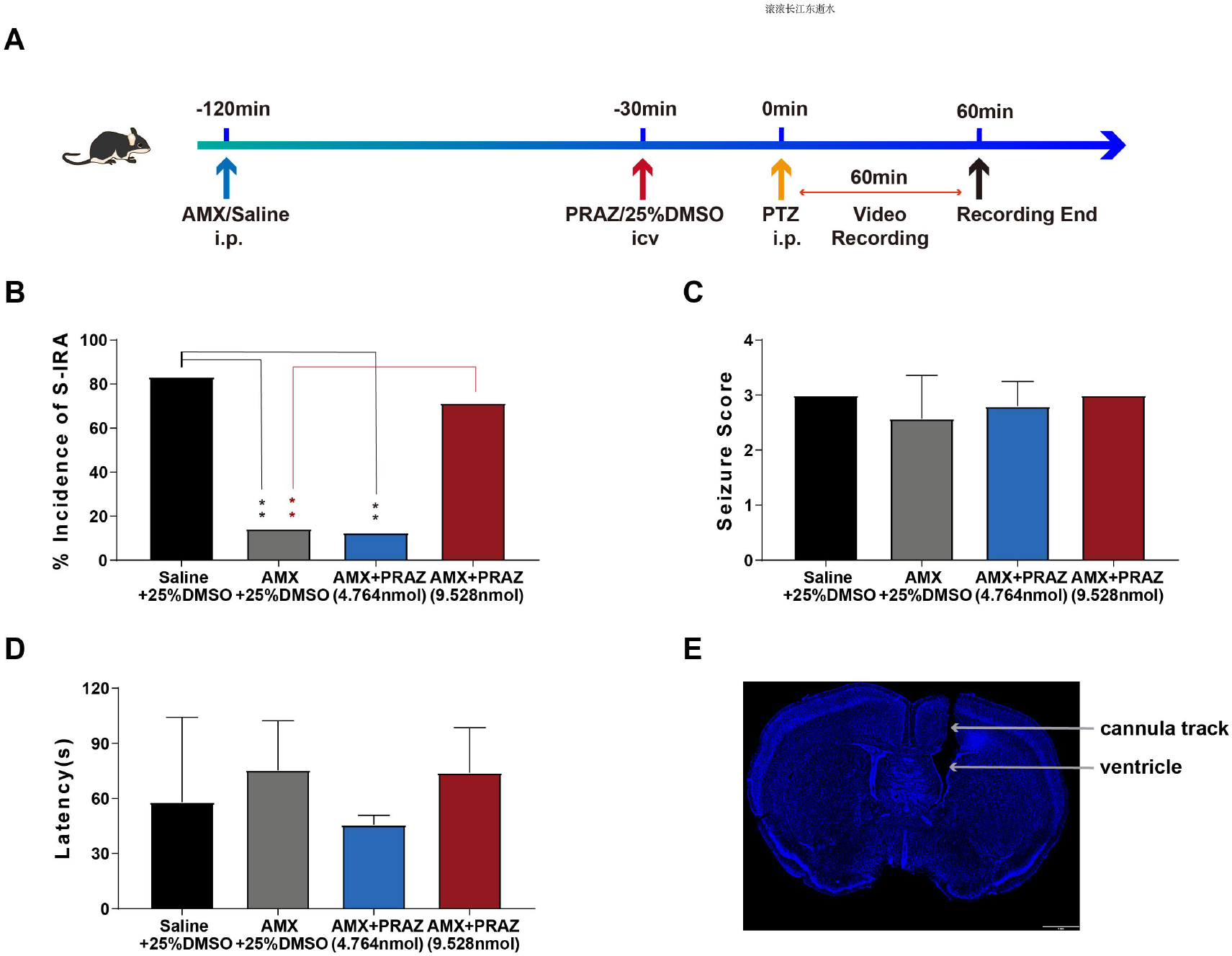
Effect of ICV prazosin on S-IRA evoked by PTZ. B. Compared to the control group (n=6) treated with vehicle (IP saline and ICV 25%DMSO), the incidence of S-IRA evoked by PTZ was suppressed in the group receiving atomoxetine (AMX) i.p. and 25%DMSO i.c.v., and the group treated with AMX i.p. and ICV prazosin (PRAZ) (4.764 nmol i.c.v.) (n=5) (**p < 0.01). The suppression of S-IRA by atomoxetine was significantly reversed by a higher dose of PRAZ (9.528 nmol ICV) (n=7) (**p<0.01). C-D. There were no significant differences of seizure scores and latencies among treated groups (p> 0.01). E. A representative histologic section indicating that vehicle or prazosin (PRAZ) were delivered successfully into the ventricle. PTZ, pentylenetetrazole; IP, intraperitoneal; ICV, intracerebroventricular; DMSO, dimethyl sulfoxide

### 4.5 ICV atomoxetine and prazosin did not attenuate PTZ-induced EEG changes

We chose the PTZ model to evaluate the effects of prazosin on EEG activity. The EEG voltage amplitude peak (power) significantly increased after PTZ injection in all three groups (Fig. 5). Meanwhile, the low frequency oscillation peak appeared before PTZ injection, and high-frequency oscillations peaked occurred post-PTZ injection in the three groups (Fig. 5, B-G). The average spectra of delta, theta and gamma bands increased after PTZ administration. These results showed that atomoxetine reduced the incidence of S-IRA without affecting seizure and EEG activity. (Fig 5, H-L).

**Figure 5.**
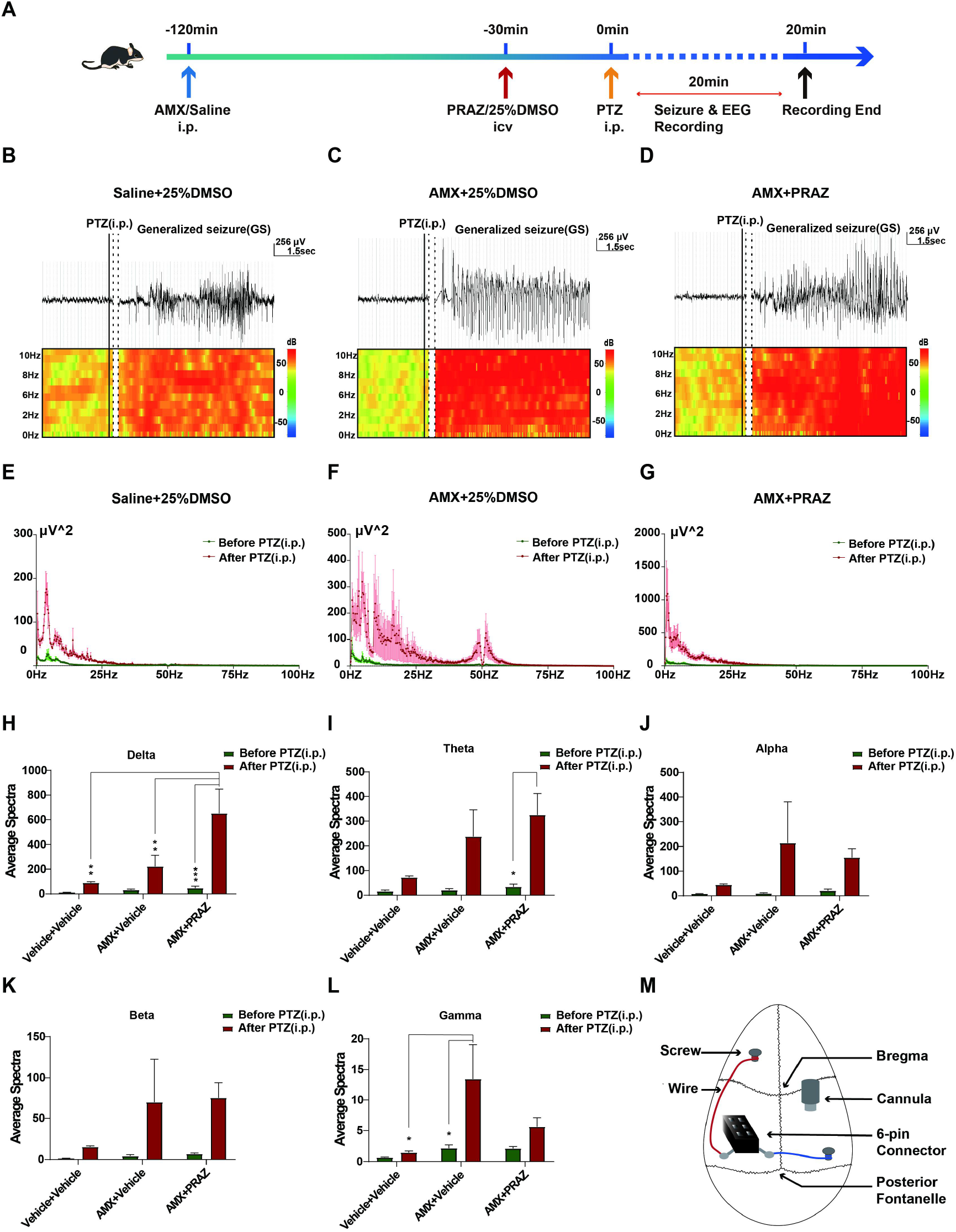
Effect of ICV prazosin on EEG in the PTZ-induced SUDEP model. A-G. Sample EEG tracings of PTZ-induced seizures in three3 different treatment groups. B-D, Changes of power and frequency oscillation peaks in the group treated with atomoxetine (AMX) 120 min and 25% DMSO i.c.v. 15 min prior to PTZ respectively, were similar to the control group. D and G. Changes of voltage amplitude and frequency oscillation peaks in the group treated with AMX + prazosin (PRAZ) (9.528 nmol, i.c.v.) were administered 120 min and 15 min prior to PTZ respectively, were not decreased compared to the control and AMX + DMSO groups. E-G. Changes of voltage amplitude peak (power) frequency oscillation peaks increased after PTZ injection in the group in which saline i.p. and 25% DMSO i.c.v. (n=3) were administered 120 min and 15 min prior to PTZ, respectively, as controls. H-L. Compared to the Vehicle + Vehicle and AMX + Vehicle groups, the average delta band spectra in the AMX+PRAZ group significantly increased (**p<0.01). I. The average theta band spectra in the AMX+PRAZ group increased after PTZ (IP) (*p < 0.05). J. There were no significant intergroup changes of average alpha band spectra. K. There were no significant intergroup differences of average beta band spectra. L. Compared to the vehicle + vehicle and AMX + vehicle groups, the average spectra of gamma band in the AMX + i.p. vehicle group increased after PTZ administration. (*p < 0.05). M. a representative illustration for the implantation of EEG electrode and ICV cannula in DBA/1 mice. PTZ, pentylenetetrazole; DMSO, dimethyl sulfoxide; i.p., intraperitoneal; i.c.v., intracerebrovascular.

## 5. DISCUSSION

Although initial progress has been made to identify the causes of SUDEP, the pathogenesis of SUDEP is still unexplored (Bateman,et al., 2010; Tomson, et al., 2008; Lhatoo, et al., 2015). Herein, our results provide new insights into the role of central norepinephrinergic neurotransmission to elucidate the cause of in SUDEP in the DBA/1 mice SUDEP model. Our data showed that the atomoxetine-mediated reduction of S-IRA was significantly reversed by administering prazosin via IP and ICV injection in AGS DBA/1 and PTZ mice SUDEP model, respectively.

### 5.1 Atomoxetine-mediated suppression of S-IRA was reversed by low-dose prazosin

Much as other investigators responded to our previous finding that atomoxetine reduced the incidence of S-IRA in a of DBA/1 murine SUDEP model by testing the role of NEα-1R to modulate S-IRA and SUDEP in lmx1bf/f/p mice with deletion of 5‐HT neurons in a maximal electroshock (MES) model (Zhang,et al., 2017; Zhao, et al., 2017; Kruse, et al., 2019), potential problems in their studies need to be considered. In the MES model, the dose of prazosin that reversed atomoxetine-mediated reduction of S-IRA was 1000 times higher than used in our DBA/1 mice SUDEP model (Kruse, et al., 2019). The higher prazosin dose may produce cardiovascular or other non-CNS side-effects that reverse the effect of atomoxetine on S-IRA in the MES model; consequently, its reversal by high-dose prazosin lacks specificity for understanding S-IRA. In addition, as the authors emphasized, the MES model of epilepsy has low fidelity to human disease, which precludes the design of clinically relevant controls.^23^

### 5.2 Atomoxetine-mediated suppression of S-IRA depends on enhanced NE bioavailability in the synaptic space by TH activity rather than TH content in the lower brainstem

Our use of two complementary murine models benefited our study design. First, we used a murine model of SUDEP evoked by acoustic stimulation to test the role of NEα-1R in modulating S-IRA as its own control. Second, we chose a model of SUDEP evoked by PTZ that recapitulates human generalized seizures to test the role of NEα-1R in modulating S-IRA. Furthermore, we explored the reversal effects of prazosin from behavioral and electrophysiologic perspectives as discussed below. Importantly, we provide the first evidence regarding brainstem TH content and activity following S-IRA, that suggests that atomoxetine-mediated inhibition of S-IRA depends on enhanced bioavailability of NE in the synaptic space and TH activity rather than TH content in the lower brainstem, suggesting that the TH activity in lower brainstem plays a key role in mediating S-IRA. However, some limitations existed in our models of SUDEP as well. Nevertheless, combining our findings of the role of NEα-1R in modulating S-IRA with those of the MES model will advance the elucidation of the pathogenesis of S-IRA and SUDEP, and inform therapeutic strategies to prevent SUDEP.

The reversal of atomoxetine-mediated reduction of S-IRA by prazosin is independent of cardiovascular function. To exclude the cardiovascular effects of prazosin on the reversal of atomoxetine-mediated suppression of S-IRA, we measured the changes of SBP, DBP, MAP and HR among different treatment groups. Prazosin used within the our dose range did not significantly change SBP, DBP, MAP and HR. However, other studies have shown that arrhythmias can cause SUDEP;^25,26^ the potential role of arrhythmias was not ruled out in our model. Prazosin is commonly used to treat hypertension, and is potentially used in epilepsy patients at risk for SUDEP; our results suggest that alternative anti-hypertensive agents should be considered for epilepsy patients. Because we previously reported that repeated generalized seizures can produce calcified cardiac lesions in DBA/1 mice,^13^ it is essential for us to clarify the role of the cardiovascular system in SUDEP in the near future.

Atomoxetine-mediated reduction of S-IRA was reversed by prazosin without affecting seizure behavior and EEG activity. We assessed the seizure score and measured the AGSZ latency and duration of clonic and/or tonic-clonic seizures to analyze seizure behavior. There were no significant intergroup differences of these parameters. Atomoxetine reduced the incidence of S-IRA without affecting EEG delta, theta and gamma activity. In addition, in our model, atomoxetine-mediated suppression of S-IRA by was reversed by prazosin in a dose-dependent manner with ceiling effects, suggesting that the central NEα-1R is a rational therapeutic target for the prevention of SUDEP. Interestingly, a recent study showed that atomoxetine-mediated suppression of S-IRA was not significantly reversed by prazosin in the same model (Zhang, et al., 2019). However, in our study, the atomoxetine-mediated suppression of S-IRA was reversed by prazosin at a dose of 0.01mg/kg (IP). The timing of prazosin administration with may have caused these discrepant results. A previous study showed that the protective effect of atomoxetine was significantly reversed by yohimbine, an NEα-2 receptor antagonist (Zhang, et al., 2019). Of course, it may be possible that both NEα-1 and NEα-2 receptors mediate SUDEP by mutual interaction.

#### Limitation to this study

A limitation of our study was that we did not identify the NEα-1R-mediated neural circuits involved in S-IRA, just as we previously had not characterized the neural circuits that suppress S-IRA evoked by either acoustic stimulation or PTZ injection in DBA/1 mice by activating serotonergic neurotransmission neurons in the dorsal raphe nucleus using optogenetic technology.^13^ Additionally, we did not establish a model of ADHD and epilepsy comorbidity to test whether the role of atomoxetine in suppression of SUDEP in ADHD and epilepsy comorbidity Nevertheless, our findings suggest that epilepsy patients receiving atomoxetine for ADHD may be receiving an additional benefit of reduced risk of SUDEP, and that the use of prazosin as an antihypertensive agent should be avoided in this patient population. Our results provide new insights into the pathogenesis of SUDEP, and suggest that atomoxetine is a promising candidate for translational research to prevent SUDEP in patients with ADHD and epilepsy comorbidity.

## Key Points

- Enhancement of central norepinephrine synthesis and norepinephrinergic neurotransmission can prevent S-IRA and SUDEP
- Atomoxetine-mediated reduction of S-IRA was significantly reversed by prazosin without affecting cardiovascular function
- The reversal of atomoxetine-induced suppression of S-IRA by prazosin has no effects on seizure behavior and EEG activity
- The activity of TH in the lower brainstem plays a key role in the pathogenesis of SUDEP
- Atomoxetine is a promising candidate for further translational research to prevent SUDEP in patients with ADHD and epilepsy

## ACKNOWLEDGEMENTS

The work was supported by the National Natural Science Foundation of China (Grant.NO: 81771403, 81974205); by the Natural Science Foundation of Zhejiang Province (LZ20H090001); by the Program of New Century 131 outstanding young talent plan top-level of Hang Zhou to HHZ; and by the National Natural Science Foundation of China (Grant.NO: 81771138) to Han Lu.

## AUTHOR CONTRIBUTIONS

HongHai Zhang designed the project; HongHai Zhang, Yi Shen and Yu Dong Zhou developed the experimental design; Yue Shen and HaiXiang Ma performed research; Han Lu and HaiTing Zhao analyzed data; Jianliang Sun and Yuan Chen interpreted data; HongHai Zhang wrote the paper.

---

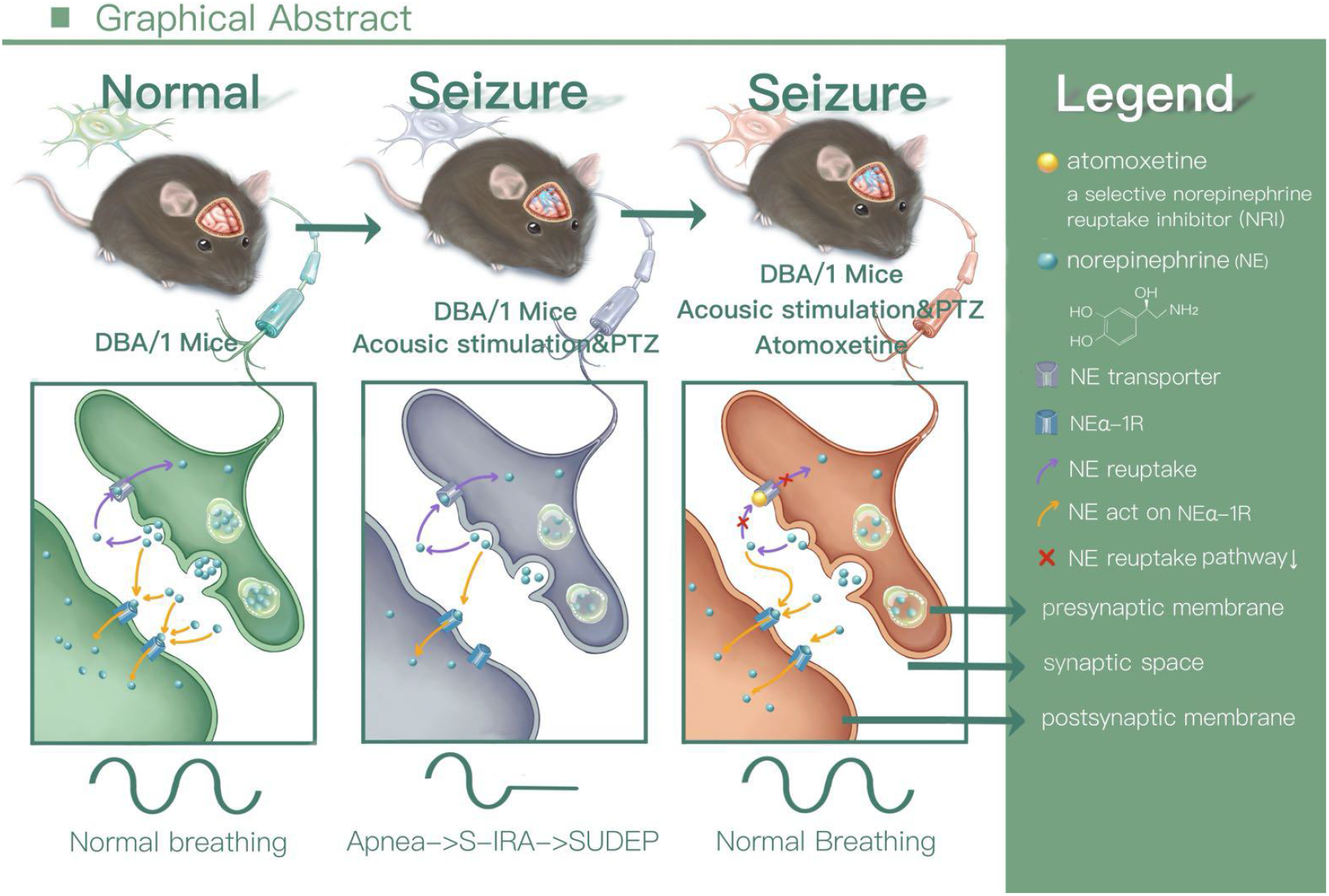

